# BGvar - a comprehensive resource for blood group immunogenetics

**DOI:** 10.1101/2021.02.04.429861

**Authors:** Mercy Rophina, Kavita Pandhare, Sudhir Jadhao, Shivashankar H. Nagaraj, Vinod Scaria

## Abstract

**Background:** Blood groups form the basis of effective and safe blood transfusion. There are about 41 well recognized human blood group systems presently known. Blood groups are molecularly determined by the presence of specific antigens on the red blood cells and are genetically determined and inherited following Mendelian principles. The lack of a comprehensive, relevant, manually compiled and genome-ready dataset of red cell antigens limited the widespread application of genomic technologies to characterise and interpret the blood group complement of an individual from genomic datasets.

**Materials and Methods:** A range of public datasets were used to systematically annotate the variation compendium for its functionality and allele frequencies across global populations. Details on phenotype or relevant clinical importance were collated from reported literature evidence.

**Results:** We have compiled the Blood Group Associated Genomic Variant Resource (BGvar), a manually curated online resource comprising all known human blood group related allelic variants including a total of 1672 ISBT approved alleles and 1552 alleles predicted and curated from literature reports. This repository includes 1606 Single Nucleotide Variations (SNVs), 270 Insertions, Deletions (InDels) and Duplications and about 1310 combination mutations corresponding to 41 human blood group systems and 2 transcription factors. This compendium also encompasses gene fusion and rearrangement events occurring in human blood group genes.

**Conclusion:** To the best of our knowledge, BGvar is a comprehensive and a user friendly resource with most relevant collation of blood group alleles in humans. BGvar is accessible online at URL: http://clingen.igib.res.in/bgvar/

## Introduction

Accurate and expedient characterization of blood group antigen profile is central to the safe and effective practice of transfusion medicine. These blood groups are defined according to the presence of specific antigens on the surface of Red Blood Cells (RBCs) with an individual’s blood type corresponding to their specific complement of RBC surface antigens^1^. Serological tests used to identify these antigens thus form the backbone of blood group characterization efforts in clinical settings.

Blood group antigens encompass a diverse range of specific genetically-determined protein sequences and oligosaccharide chains. With the exception of de-novo events, blood groups thus exhibit a Mendelian pattern of inheritance. Human blood group systems are often regulated by a single gene locus or two or by more closely linked homologous genes. There exists about 360 human blood group antigens, 322 of which belong to 36 well recognized blood group systems^2^.

Virtually all genes involved in encoding these human blood group antigens have been identified and mapped^3^. One of the earliest reviews elaborating the genetic basis of the human ABO blood group system extensively explained the types of variations and their functional impacts^4,5^. RH is yet another major human blood group system whose molecular characterization has been pervasively studied^6,7,8^. The accurate characterization of these blood groups is important both in the context of preventing transfusion-related hemolytic reactions, and in research efforts aimed at understanding the pathogenesis of a number of hematological disorders^1,9,10,11^.

The advent of high throughput genomic modalities including array based genotyping and Next Generation Sequencing (NGS) technologies has revolutionized the application of genomics in clinical settings. In recent years, these technologies have enabled the characterization of potentially clinically-relevant genetic variations, among individuals. Such characterization is potentially relevant for the diagnosis of genetic conditions for pharmacogenomics, for optimizing therapeutic dosing strategies, and for minimizing adverse events while simultaneously identifying genetic correlates of a number of clinical traits and conditions. With its efficacy in overcoming most of the limitations borne by serological and molecular diagnostics, it has now become the future choice in blood group genotyping strategies. One of the major and significant advantage of NGS based blood typing is the ability to determine the sequence of the entire gene coding for the red blood cell/platelet antigen, thereby uncovering the role of novel variants in human blood group systems.^12,13,14,15^ Extensive blood typing procedures are also commonly employed to minimize transfusion reactions in clinical settings^16^. The need for such extensive characterization is also highlighted by a recent annual report of the Food and Drug Administration (FDA) regarding transfusion associated fatalities, which emphasizes that ∼7% of Hemolytic Transfusion Reactions (HTRs) observed in the United States from 2013 - 2017 were due to ABO incompatibilities whereas, 11% of these reactions were due to non-ABO incompatibilities^17^.

Despite the promise of these personalized blood typing strategies, the widespread use of genomic technologies in transfusion medicine is presently hampered by the lack of a systematic and relevant compendium of alleles associated with blood group antigens that can aid the clinical interpretation of genetic variants based upon the results of high throughput experiments. One of the initiatives by the National Center for Blood Group Genomics (NCBGG) has maintained a compilation of about 148 RHCE alleles along with their relevant information like ancestry, prevalence and variation IDs^18^. To that end, we created BGvar, a comprehensive catalog of genetic variants associated with human blood group systems. To the best of our knowledge, BGvar is the most comprehensive and up-to-date compendium of human blood group alleles. The resource is accessible online at URL: http://clingen.igib.res.in/bgvar/

## Materials and Methods

### Datasets

Human blood group variant information was compiled from literature sources and public documents. The compiled variants were additionally checked in public resources and appropriate identifiers were mapped. Blood group antigens and allele data was manually collated from resources including the International Society of Blood Transfusion (ISBT)^2^, The Blood Group Antigen Gene Mutation Database (BGMUT)^19^, Erythrogene^20^, The Human RhesusBase^21^, Blood Antigens^22^,^23^ and literature references including *The Blood group antigen factsbook*^*24*^ and *Blood groups and Red cell antigens*^*25*^. ISBT has systematically developed and maintained a repository of blood antigen information corresponding to 41 human blood groups and 2 transcription factors. BGMUT, a now inactive online resource, comprises a manually curated compilation of allelic variations in various human blood group related genes accounting for about 1251 alleles from 41 genic loci. Erythrogene is an online search engine which comprises multiethnic human blood group related alleles predicted from 1000 Genomes Project in addition to known and recognized alleles. RhesusBase is a recent resource that exclusively encompasses Rhesus and Colton blood group antigen information. Blood Antigens resource, in addition to compilation of molecular changes occurring in Red Blood Cells and Platelets, also provides an automated blood antigen - typing algorithm from genome sequencing data (BloodTyper). A schematic representation of the data compilation process is summarized in **Figure 1**.

**Figure 1.**
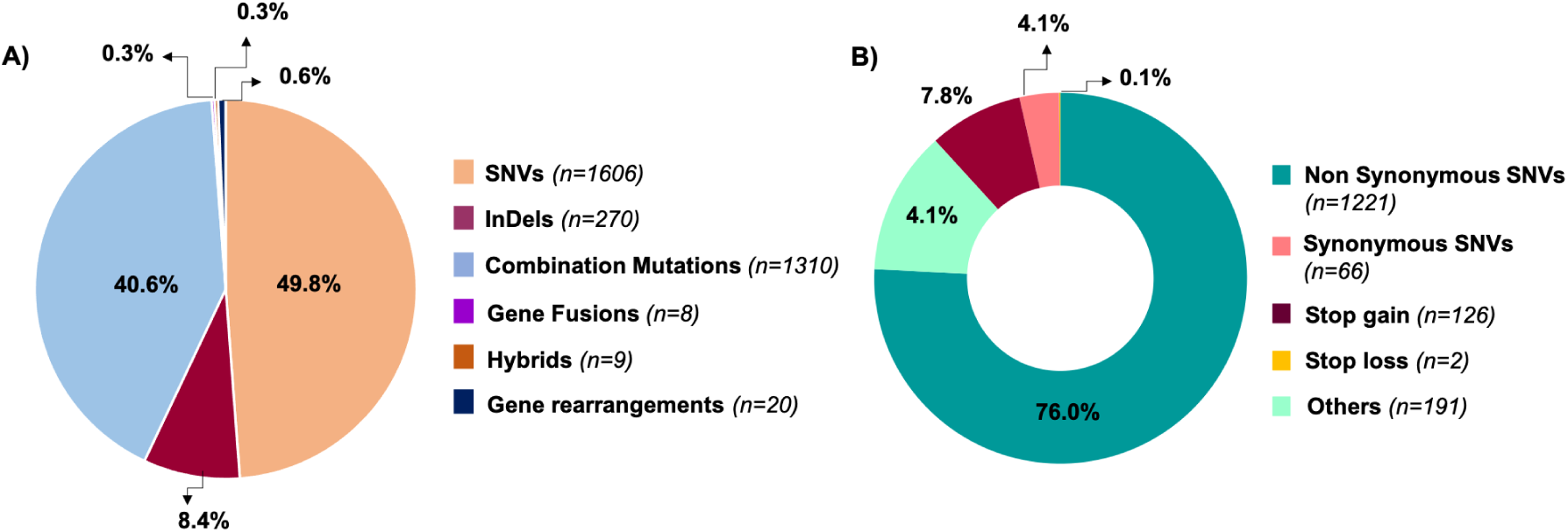
Schematic summarising the data curation, database mapping, variant annotation and database/interface development of BGvar

### Compilation and annotation of genetic variants

All variants were first individually compiled to a master sheet and identifiers were mapped. Blood group IDs and gene names were obtained for each variant. Phenotype and allele names (as per ISBT nomenclature) were also collected. Genomic loci corresponding to these variants were obtained from the RefSeq database for both GRCh37/hg19 and GRCh38/hg38 human genome assemblies. Sequence variant descriptions for DNA, RNA and protein levels (according to HGVS nomenclature)^26^ were added and verified using Mutalyzer^27^,^28^. Variant annotation was performed using Annotate Variation (ANNOVAR)^29^ integrating a number of public databases and computational tools. The allelic frequency of each variant was retrieved from global population datasets including the 1000 Genomes project^30^, Exome Sequencing Project (ESP6500), Exome Aggregation Consortium (ExAC v.0.3)^31^, Genome Aggregation Database (gnomAD v2.1.1 Exomes) and The Greater Middle East (GME) Variome project^32^. Comprehensive list of datasets used for variant annotation is summarized in **Supplementary Table 1**. The clinical significance of the variants in the compendium was evaluated individually through manual literature searches and curation. Comparison of BGvar features with other existing databases is summarized in **Supplementary table 2**.

**Table 1a.**
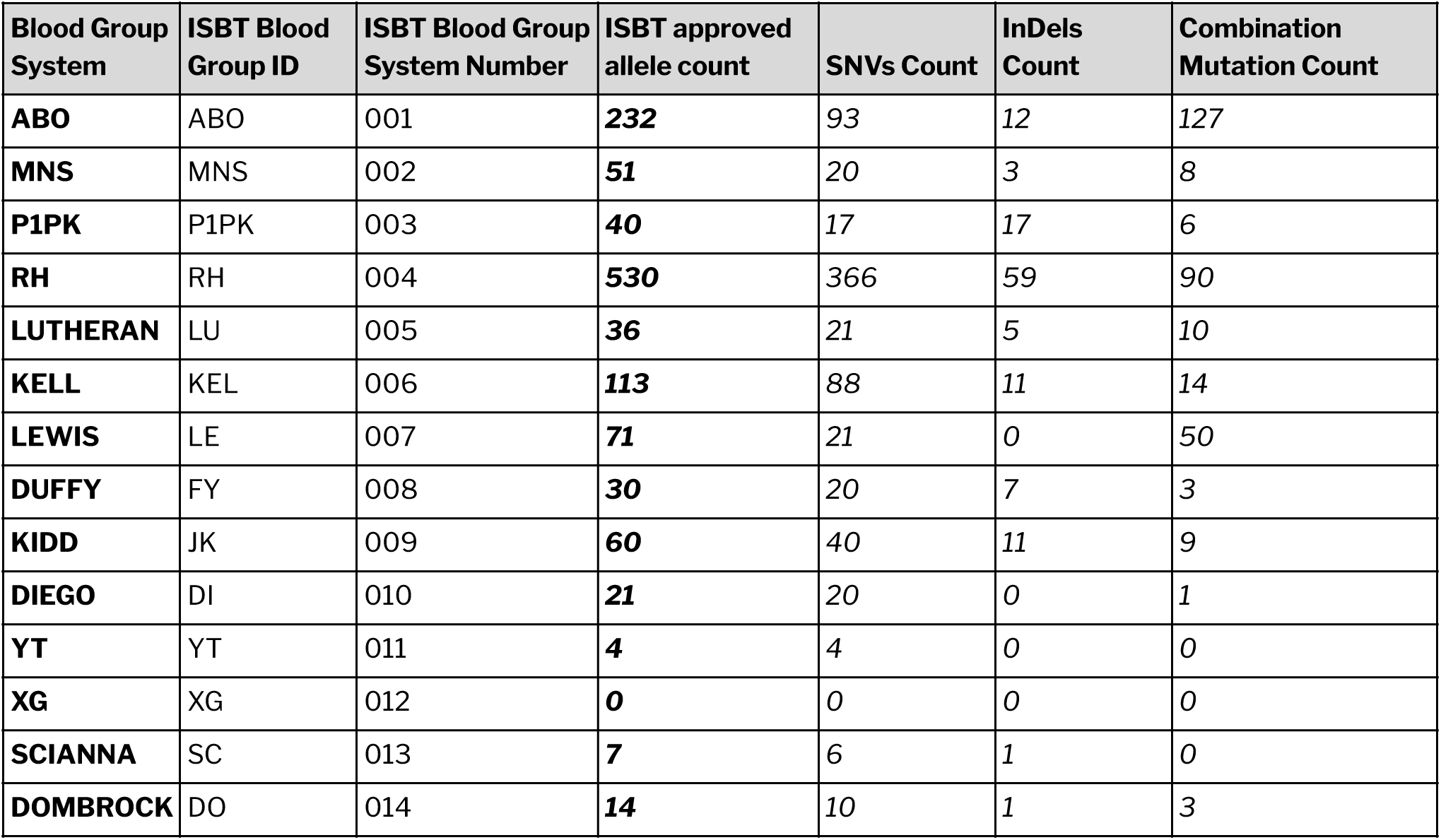
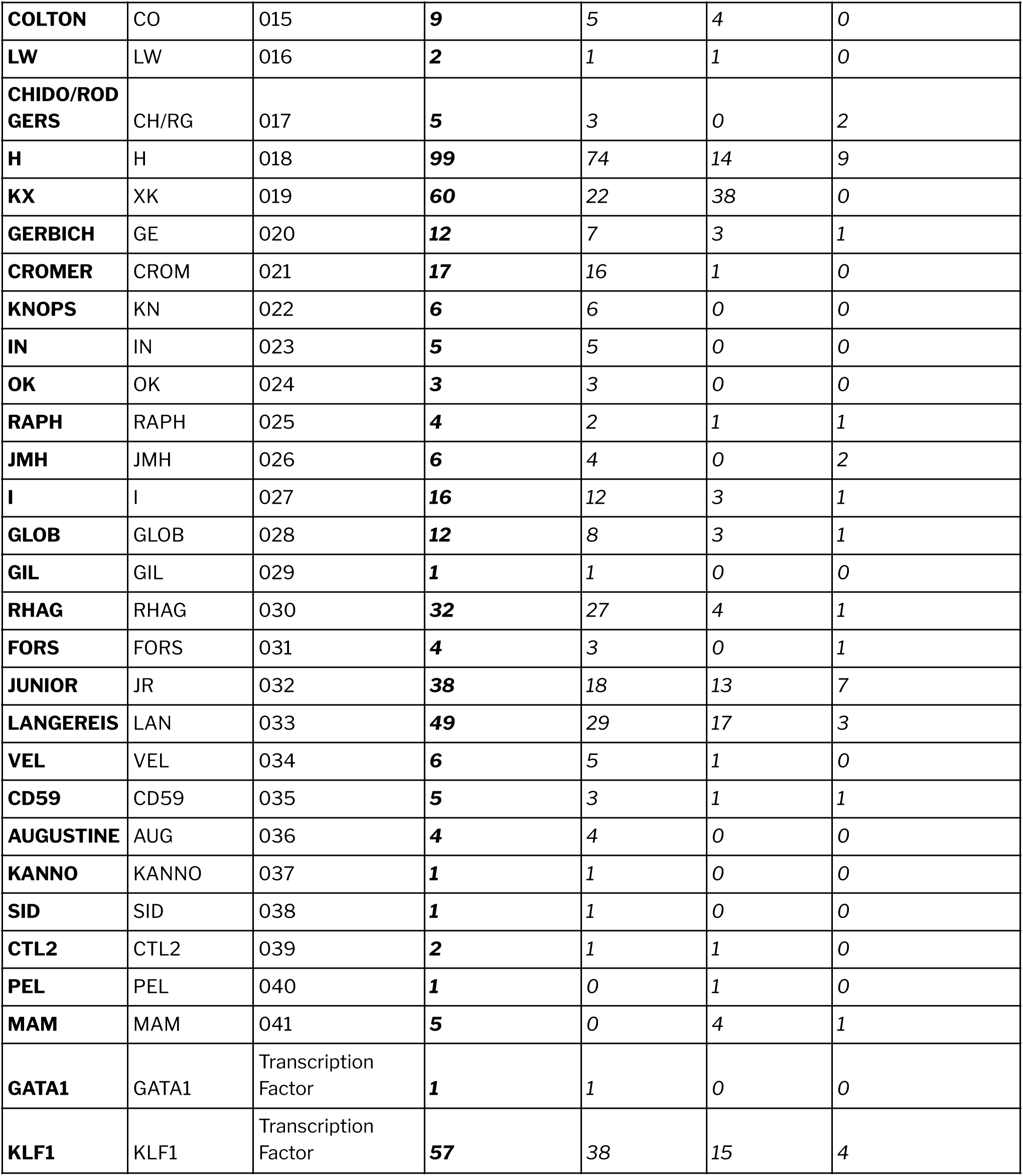
Summary of ISBT approved human blood group alleles and their classifications

**Table 1b.** Summary of human blood group alleles predicted from the 1000 Genomes Project and curated from literature evidence

| Blood Group System | ISBT Blood Group ID | ISBT Blood Group System | Predicted and curated allele | SNVs Count | InDels Count | Combination Mutation Count |
| --- | --- | --- | --- | --- | --- | --- |

**Table 1b.** Summary of blood group alleles in each blood group system predicted from the 1000 Genomes Project
|  |  | Number | count |  |  |  |
| --- | --- | --- | --- | --- | --- | --- |
| **ABO** | ABO | 001 | **72** | 38 | 6 | 28 |
| **MNS** | MNS | 002 | **41** | 12 | 0 | 29 |
| **P1PK** | P1PK | 003 | **33** | 4 | 0 | 29 |
| **RH** | RH | 004 | **104** | 13 | 0 | 91 |
| **LUTHERAN** | LU | 005 | **80** | 32 | 0 | 48 |
| **KELL** | KEL | 006 | **71** | 32 | 0 | 39 |
| **LEWIS** | LE | 007 | **72** | 22 | 0 | 50 |
| **DUFFY** | FY | 008 | **13** | 2 | 0 | 11 |
| **KIDD** | JK | 009 | **27** | 0 | 0 | 27 |
| **DIEGO** | DI | 010 | **63** | 44 | 0 | 19 |
| **YT** | YT | 011 | **29** | 16 | 0 | 13 |
| **XG** | XG | 012 | **5** | 5 | 0 | 0 |
| **SCIANNA** | SC | 013 | **44** | 25 | 0 | 19 |
| **DOMBROCK** | DO | 014 | **28** | 8 | 0 | 20 |
| **COLTON** | CO | 015 | **22** | 20 | 0 | 2 |
| **LW** | LW | 016 | **11** | 10 | 0 | 1 |
| **CHIDO/RODGERS** | CH/RG | 017 | **44** | 17 | 0 | 27 |
| **H** | H | 018 | **79** | 34 | 8 | 37 |
| **KX** | XK | 019 | **2** | 2 | 0 | 0 |
| **GERBICH** | GE | 020 | **15** | 4 | 0 | 11 |
| **CROMER** | CROM | 021 | **16** | 16 | 0 | 0 |
| **KNOPS** | KN | 022 | **273** | 4 | 0 | 269 |
| **IN** | IN | 023 | **17** | 14 | 0 | 3 |
| **OK** | OK | 024 | **19** | 7 | 0 | 12 |
| **RAPH** | RAPH | 025 | **17** | 8 | 0 | 9 |
| **JMH** | JMH | 026 | **34** | 13 | 0 | 21 |
| **I** | I | 027 | **32** | 30 | 0 | 2 |
| **GLOB** | GLOB | 028 | **4** | 4 | 0 | 0 |
| **GIL** | GIL | 029 | **9** | 1 | 0 | 8 |
| **RHAG** | RHAG | 030 | **27** | 24 | 0 | 3 |
| **FORS** | FORS | 031 | **63** | 20 | 0 | 43 |
| **JUNIOR** | JR | 032 | **60** | 35 | 0 | 25 |
| **LANGEREIS** | LAN | 033 | **48** | 5 | 0 | 43 |
| **VEL** | VEL | 034 | **14** | 12 | 2 | 0 |
| **CD59** | CD59 | 035 | **5** | 4 | 0 | 1 |
| **AUGUSTINE** | AUG | 036 | **18** | 18 | 0 | 0 |
| **KANNO** | KANNO | 037 | **0** | 0 | 0 | 0 |
| **SID** | SID | 038 | **0** | 0 | 0 | 0 |
| **CTL2** | CTL2 | 039 | **0** | 0 | 0 | 0 |
| **PEL** | PEL | 040 | **0** | 0 | 0 | 0 |
| **MAM** | MAM | 041 | **0** | 0 | 0 | 0 |
| **GATA1** | GATA1 | Transcription<br>Factor | **16** | 14 | 1 | 1 |
| **KLF1** | KLF1 | Transcription<br>Factor | **25** | 11 | 1 | 13 |

### Database architecture

The variant data and corresponding annotations were transformed to JavaScript Object Notation format and ported onto MongoDB 3.4.1. The user-friendly web interface for querying the database was coded in PHP 7.0, AngularJS, HTML, Bootstrap 4 and CSS. The web server was configured in Apache HTTP server. The Highcharts javascript library was used for the graphical representation of variant data.

## Results

### Compendium of blood group variants

BGvar comprises 3224 human blood group related variants. This compilation precisely includes 1672 ISBT approved alleles and 1552 alleles predicted from 1000 Genomes Project and compiled from literature. The repository holds 1606 Single Nucleotide Variations (SNVs), 270 Insertions, Deletions (InDels) and duplications and about 1310 combination mutations corresponding to 41 human blood group systems and 2 transcription factors. Genetic variations due to events like gene fusion and rearrangements were also included. Of the total SNVs, 1016 were found to be non synonymous, 66 were synonymous, 126 and 2 were stop gain and stop loss variants respectively. Detailed summary of blood group related variants compiled in the resource is provided in **Table 1a and 1b. Figure 2** illustrates the category and function classification of the compiled variants.

**Figure 2.**
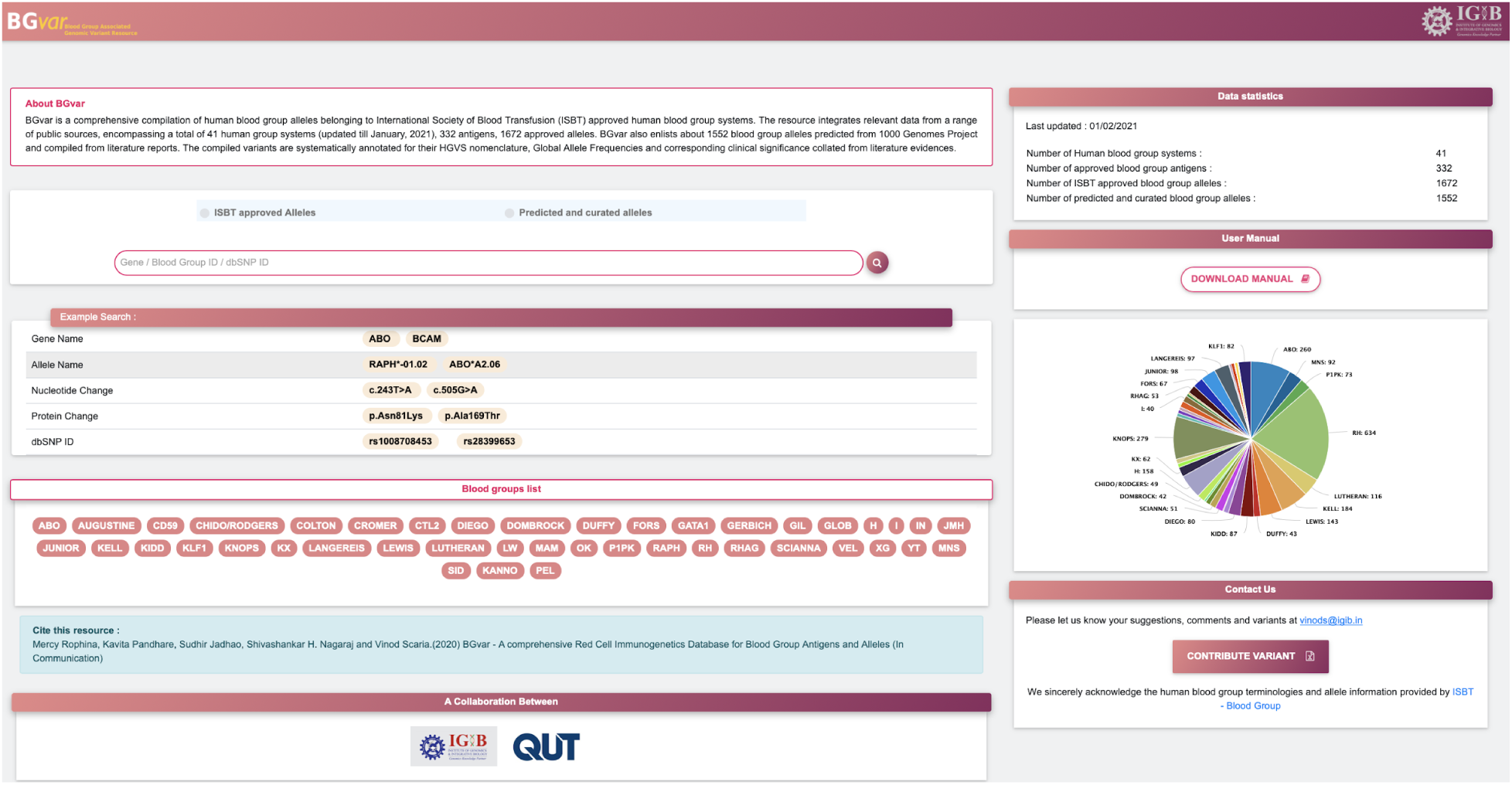
Illustration of categories and functional classifications of variants compiled in the resource

### Database features and navigation

BGvar has been designed to have a user friendly interface. Blood group systems in this database are organized on the homepage for rapid access to relevant information. In addition, the search interface enables the user to query the database based on gene name, genetic variant ID (rsID), nucleotide change, protein change and ISBT allele name. This database navigates through 3 major sections: (i) Basic information regarding the blood group system, providing a layman’s summary of the blood group and its history; (ii) A blood group variant list with summary information and (iii) A detailed description of the variants associated with the blood group. Users are also provided with an option to search for ISBT approved alleles and other predicted and curated alleles separately. **Figure 3a and 3b** illustrates various sections of result display of the resource.

**Figure 3.**
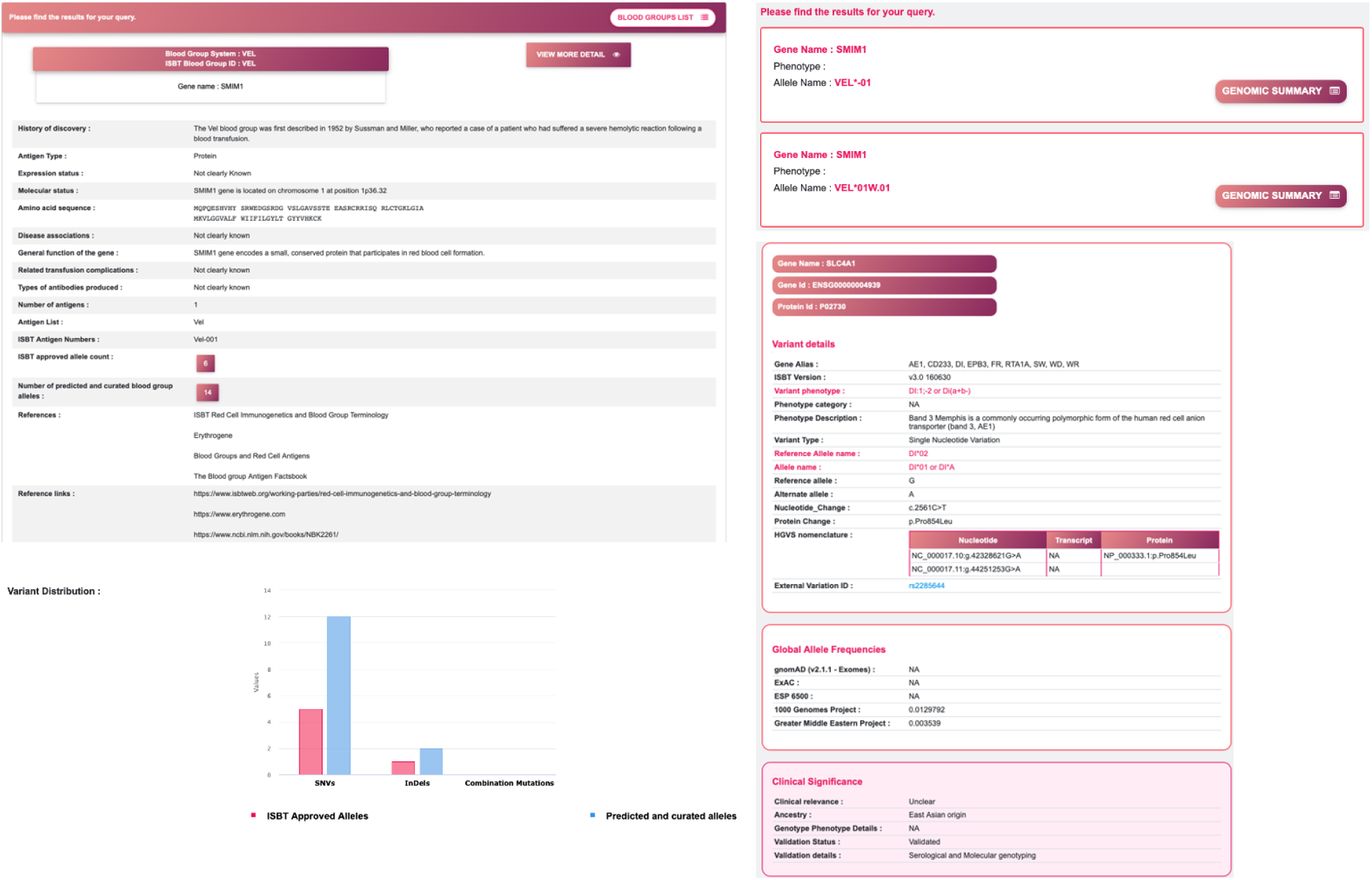
Illustration of query search and sections of result display in BGvar. Homepage of the web resource with query search examples and user contribution options *(top)*. Result display sections which provide brief information pertaining to the blood group system, list of alleles known and reported and the complete genomic summary of a selected variant *(bottom)*.

### Blood group information

BGvar provides a brief summary pertaining to each blood group system. The section briefly details the history of the blood group system including its discovery, antigen expression patterns, molecular details and information regarding gene functionality. This is followed by the amino acid sequence of the protein, and information regarding reported disease associations, if any. Brief summary on the list of ISBT approved antigens and alleles pertaining to each blood group system along with the counts of alleles predicted from the 1000 Genomes project data and curated from literature sources are also provided. Additionally, this section details the clinical correlates or transfusion complications, if any, derived from literature.

### Genetic Variant Information

This section details the genetic variants associated with each blood group. These include basic details pertaining to these variants, such as allele names, allele phenotypes, genomic loci, cDNA sequences and protein changes conforming to the HGVS nomenclature. This section also contains information regarding the corresponding global population frequencies of a given variant in major global datasets including the gnomAD (version 2.1.1 exomes), ExAC, 1000 Genomes Project, Greater Middle East and ESP6500. This page also details any known information regarding the clinical significance of the variant and phenotypic or genotypic descriptions thereof.

### Community participation in variant curation/annotation

As blood group systems and genetic variants associated with blood groups are being continuously discovered, it is imperative that reference databases be up-to-date. BGvar therefore invites the community to contribute and enrich the utility this resource. BGvar provides an option for users to contribute any blood group related variant discoveries to the repository using a standard template. A detailed user manual to guide users in accessing the database is also provided.

## Discussions

Genome-scale technologies have significantly impacted many clinical fields, and have promised to pave the way for precision medicine. Such genomic technologies are anticipated to significantly impact transfusion medicine, by facilitating a more precise understanding of the genetic repertoire of variants associated with blood group systems^33,34^. Such an approach would have implications for the identification of rare blood groups, banking of blood and blood products and safer transfusion. As the numbers of new blood antigens being identified and reported continues to rise, a comprehensive compendium has the potential to serve as a reliable reference for transfusion research. BGvar is designed to fit this niche by compiling data pertaining to 3224 antigens belonging to 41 human blood group systems. We publish this resource in the hopes that it will be widely used by the community and will form a central resource for blood group immunogenetics.

## Acknowledgements

MR curated the data for the database. KP designed the database. SJ and SHN helped in fact checking. MR and VS contributed in writing the manuscript. VS conceived and designed the project. All authors approved the final manuscript. Authors acknowledge help, support and constructive comments from Mukta Poojary, Bani Jolly and Ambily Sivadas. Authors acknowledge funding from CSIR India. The funders had no role in the preparation of the manuscript or decision to publish.

---

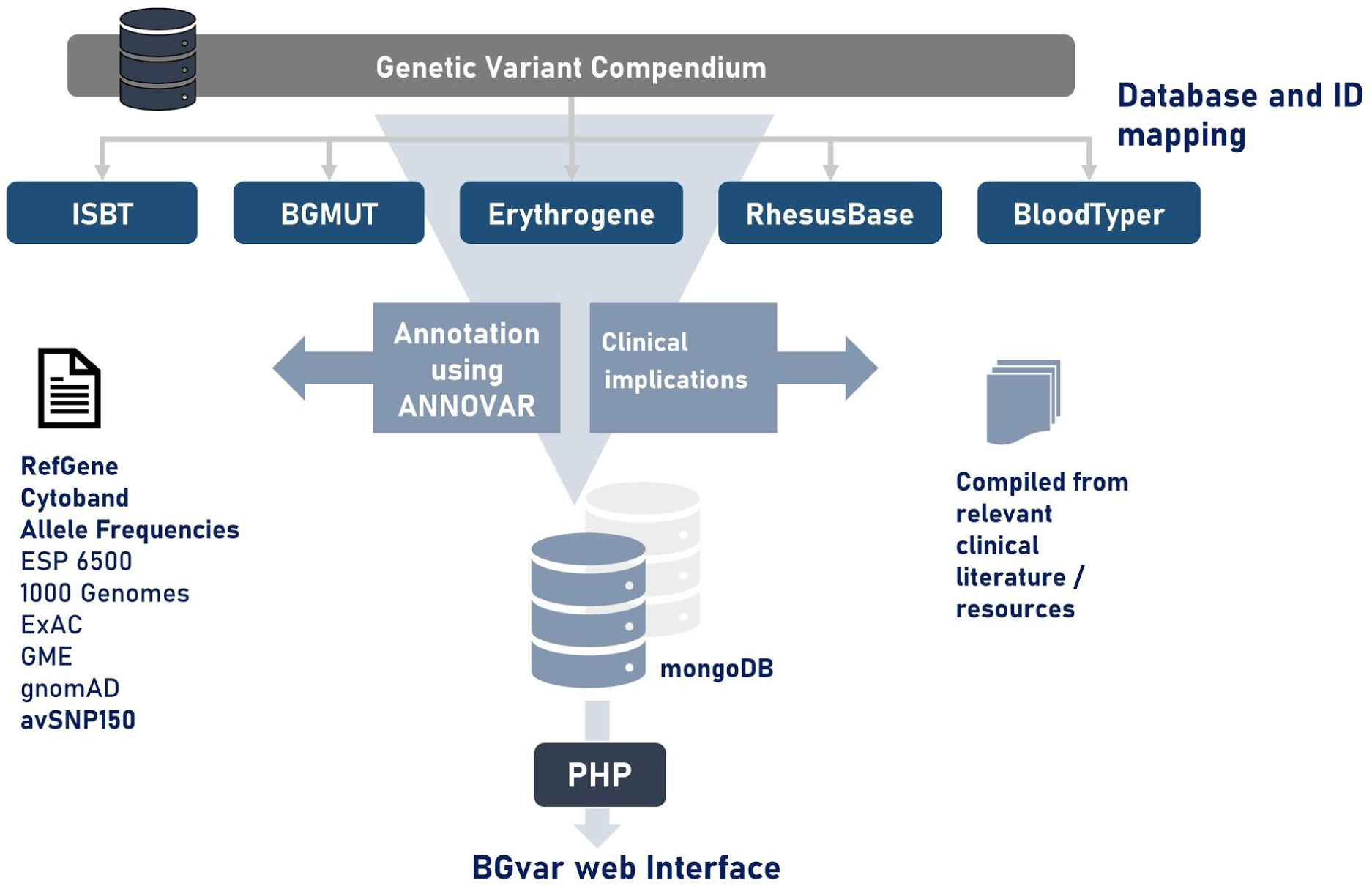

