## Supplementary Data for "BGvar - a comprehensive resource for blood group immunogenetics"

**Supplementary Table 1.** Comparison of BGvar with other existing databases.

**Supplementary Table 2.** Summary of datasets used for variant annotation.

|  | **BGMUT** | **ISBT** | **Erythrogene** | **RhesusBase** | **BloodTyper** | **BGvar** |
| --- | --- | --- | --- | --- | --- | --- |
| ***Blood group system overview*** | No | Yes | No | No | No | Yes |
| ***Date of last update*** | Inactive | 2020 | 27 Nov 2017 | 18 Mar, 2020 | 08 Sept, 2020 | Jan, 2021 |
| ***Blood group allele information*** | Manual curation of blood group related alleles from literature sources | Provides standard blood group allele nomenclature and phenotype information | Enlists ISBT approved alleles and blood group related alleles predicted from 1000 Genomes project | Exclusively comprises ISBT approved RH and CO blood group alleles | Provides list of ISBT approved alleles | Enlists ISBT approved as well as alleles predicted from 1000 Genomes dataset |
| ***Variant annotation*** | No | Provides the nucleotide and the amino acid change | Provides the nucleotide and the amino acid change | Provides the nucleotide and the amino acid change | Provides the nucleotide and the amino acid change | Categorizes the genetic variations responsible for the blood group phenotype along with extensive functional annotation |
| ***HGVS nomenclature*** | No | No | No | No | No | Yes  (Genomic, Transcript and Protein level) |
| ***Global allele frequencies*** | Brief mention on the prevalence (if any) | Reports the prevalence (if any) | Reports the frequency of the allele from 1000 Genomes data | No | Reports data from Exon Variant Server | Provides the allele frequency calculated from global datasets including **ESP6500, EXAC, 1000 Genomes, gnomAD and GME** |
| ***Ancestry details*** | Yes | Yes | No | No | No | Yes  (Manual curation from corresponding literature sources) |
| ***Phenotype/Genotype details*** | No | No | No | No | No | Yes  (Manual compilation of information provided from the reported case studies - if any) |
| ***Clinical significance*** | No | No | No | No | Yes  General description on the hemolytic transfusion complication status | Yes (Manual curation from reported literature - if any) |
| ***Web interface*** | No (Inactive -Data available) | Yes | Yes | Yes | Yes | Yes |
| ***Search interface*** | No | No | Yes | No | Yes | Yes |
| ***Link to resources*** | No | No | No | No | Yes | Yes |
| ***User contribution options*** | Not active | Generates the framework for inclusion of new discoveries | No | No | No | Yes  A user friendly method of sharing new allele discoveries to the repository |

Table 1 : Comparison of BGvar with other existing databases

| **Gene based annotations** | |
| --- | --- |
| ***Dataset name*** | ***Description*** |
| RefGene | Provides gene and transcript level annotations. Amino acid changes for nonsynonymous variants are also shown. |
| CytoBand | Provides precise genomic location of the gene |
| **Filter based annotations** | |
| ***Dataset name*** | ***Description*** |
| Esp6500 | Provides allele frequency of the variants from NHLBI-ESP 6500 |
| 1000g (Aug-2015) | Provides allele frequency of the variants from The 1000 Genomes Project |
| Exac (v3.0) | Provides allele frequency of the variants from The Exome Aggregation Consortium (version 3) |
| GME | Provides allele frequency of the variants from The Greater Middle East genetic variant compilation |
| gnomAD (2.1.1 exomes) | Provides allele frequency of the variants from the Genome Aggregation Database (gnomAD; version 2.1.1 - Exome data) |
| avsnp150 | Precisely maps the identifiers for the genetic variants provided in dbSNP domain |

**Supplementary Table 1.** Brief description of various datasets used for variant annotation
